# Human Mucosal Associated Invariant T cell proliferation is dependent on a MYC-SLC7A5-Glycolysis metabolic axis

**DOI:** 10.1101/2022.01.17.476571

**Authors:** Nidhi Kedia-Mehta, Marta M. Pisarska, Christina Rollings, Chloe O’Neill, Conor De Barra, Cathriona Foley, Nicole AW. Wood, Neil Wrigley-Kelly, Natacha Veerapen, Gurdyal Besra, Ronan Bergin, Nicholas Jones, Donal O’Shea, Linda V. Sinclair, Andrew E. Hogan

## Abstract

Mucosal Associated Invariant T (MAIT) cells are an abundant population of innate T cells which recognise bacterial ligands presented by the MHC class-I like molecule MR1. MAIT cells play a key role in host protection against bacterial and viral pathogens. Upon activation MAIT cells undergo proliferative expansion and increased production of effector molecules such as cytokines. The molecular and metabolic mechanisms controlling MAIT cell effector functions are still emerging. In this study, we found that expression of the key metabolism regulator and transcription factor MYC is upregulated in MAIT cells upon immune stimulation. Using quantitative mass spectrometry, we identified the activation of two MYC controlled metabolic pathways; amino acid transport and glycolysis, both of which are critical for MAIT cell proliferation. Finally, we show that MYC expression in response to immune activation is diminished in MAIT cells isolated from people with obesity, resulting in defective MAIT cell proliferation and functional responses. Collectively our data details for the first time the importance of MYC regulated metabolism for MAIT cell proliferation, and provides additional insight into the molecular defects underpinning functional failings of MAIT cells in obesity.

**Graphical Abstract:** 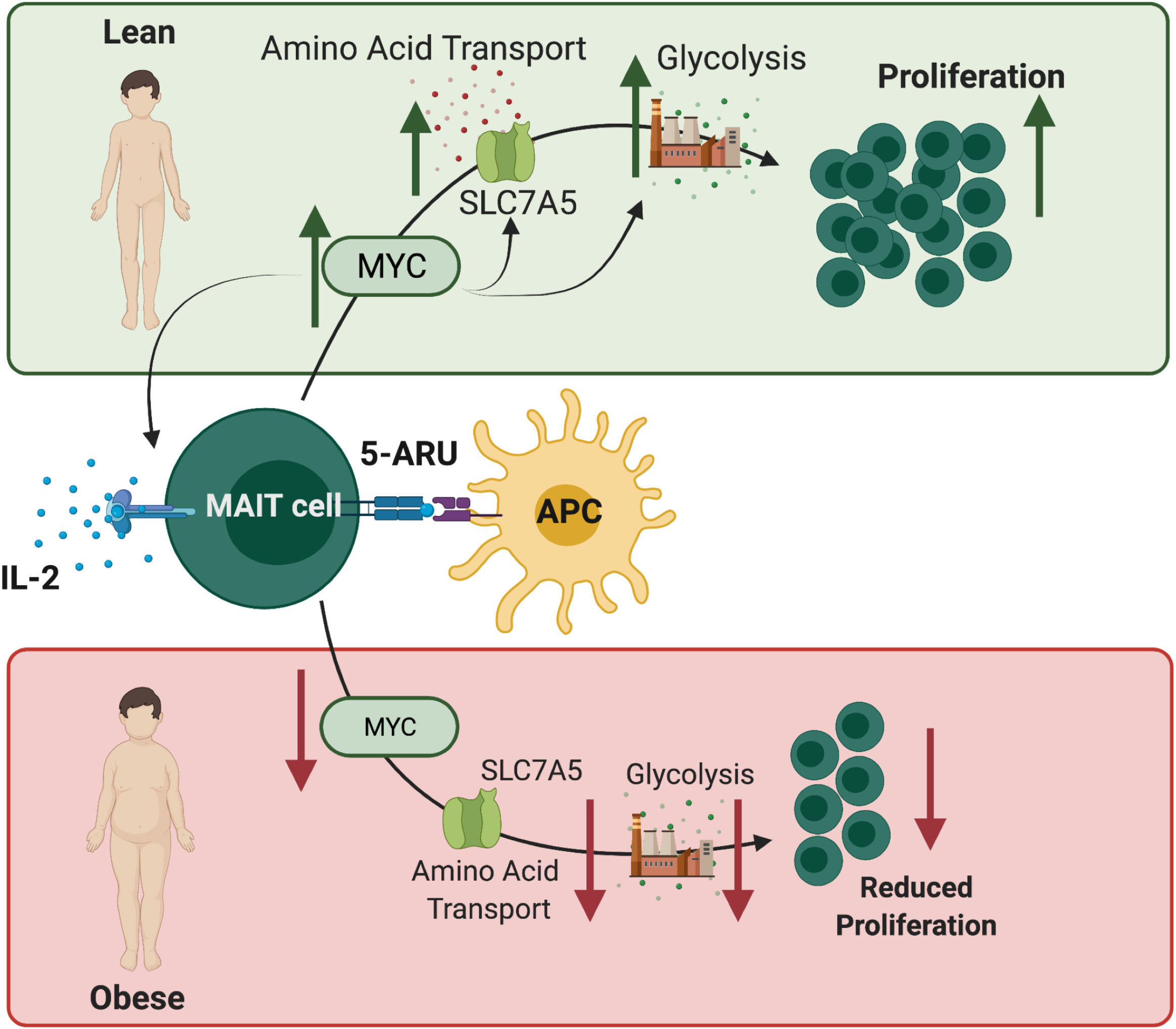

## Introduction

MAIT cells are a population of non-MHC restricted T cells that are important in the immune defence against bacterial and viral infections^1, 2, 3, 4, 5^. MAIT cells are early responding T cells that are capable of rapidly producing multiple cytokines upon activation such as IFN-g, TNF and IL-17^1, 6^. MAIT cells are activated when their invariant TCR recognise bacterial derivatives presented on the MHC like molecule MR1^5, 6^. MAIT cells can also be activated in a T cell receptor independent manner, via cytokine stimulation^7, 8^. Dysregulated MAIT cell cytokine profiles have been reported in several diseases including obesity, arthritis and viral infection^9, 10, 11, 12, 13^.

One key functional response of MAIT cells is their ability to rapidly proliferate upon activation and has been demonstrated both *in vitro* and *in vivo*^14, 15, 16^. This increased immune-signal driven proliferation relies upon metabolic reprogramming to provide the significant amounts of energy and biosynthetic intermediates needed to support the rapid generation of new cells^17^. In conventional T cells, metabolic reprogramming in response to activation is controlled by the transcription factor MYC, which acts as a metabolic master regulator^18^. MYC expression is rapidly induced after T cell receptor (TCR) stimulation, and is sustained by amino acid availability and IL-2 signalling^19, 20,21^. In particular the regulated expression of the amino acid transporter SLC7A5, which is under the control of MYC, forms a forward feeding loop where amino acid transport via SLC7A5 sustains MYC protein expression^19^. Whether MYC acts as a metabolic regulator in MAIT cells is currently unknown. We have previously demonstrated that MAIT cell cytokine production is dependent on glycolytic metabolism, however it is unclear what regulates and fuels MAIT cell proliferation^22^. To address these unknowns, we have interrogated the molecular and metabolic requirements MAIT cell proliferation.

Using quantitative mass spectrometry, we identify a robust upregulation of MYC expression and MYC controlled metabolic pathways in stimulated MAIT cells. We show that MYC-regulated metabolic pathways (including amino acid transport and glycolysis) are essential for human MAIT cell proliferation. Finally, we show that obesity is associated with defective MAIT cell proliferation, due to a defective MYC-SLC7A5-Glycolysis metabolic axis. Collectively our data demonstrate that MYC acts as a metabolic regulator in TCR and cytokine stimulated MAIT cells and that this is essential for proliferation, and provides further molecular insight into obesity related defects in human MAIT cells.

## RESULTS

### Quantitative mass spectrometry analysis reveals MAIT cell proteomic remodelling upon activation

To investigate the major pathways regulating MAIT cell activation, we performed quantitative mass spectrometry on IL-2 expanded MAIT cells (MAIT^IL-2^ cells) before and after 18 hours of stimulation (anti-TCR/CD28 and IL-12/IL-18; MAIT^STIM^). We identified over 6800 proteins in both IL-2 maintained and stimulated MAIT cells, and have estimated the absolute protein copy number per cell using the “proteomic ruler” method which uses the mass spectrometry signal of histones as an internal standard^23^. As has previously been described in conventional effector T cells, only a very small proportion of highly abundant proteins account for the majority of cellular mass^19, 24, 25^; in the context of these human MAIT cells, we find that expression of 7 proteins contribute 25% of the total mass, and expression abundance of only ~ 350 proteins is the basis of 75 % of the total cellular mass (Figure 1A.) Upon activation, we observed a significant remodelling of the MAIT^IL-2^ cell proteome (Figure 1B), which was accompanied by an increase in protein mass and cell size (Figure 1C-D). Of note, the expression of the most abundant proteins did not change with regard to stimulation, remaining proportional to the levels expressed basally in MAIT^IL-2^ cells. Signature effector proteins such as Interferon gamma (IFNG) and Granzyme B (GRZB) were amongst the most highly increased proteins upon stimulation. We were also able to identify expression of many core MAIT cell proteins including IL-18R1 (which strongly increased expression in response to stimulation), KLRB1 (moderate increase), DPPIV (moderately decreased), and TRAV1-2 (expression is decreased upon stimulation) (Figure 1E). In total, over 1400 proteins were increased more than 1.5-fold upon activation (Figure 1F). Pathway analysis highlighted that the increased proteins were enriched in multiple processes including a significant proportion involved in governing cellular metabolism (Figure 1G).

**Figure 1.**
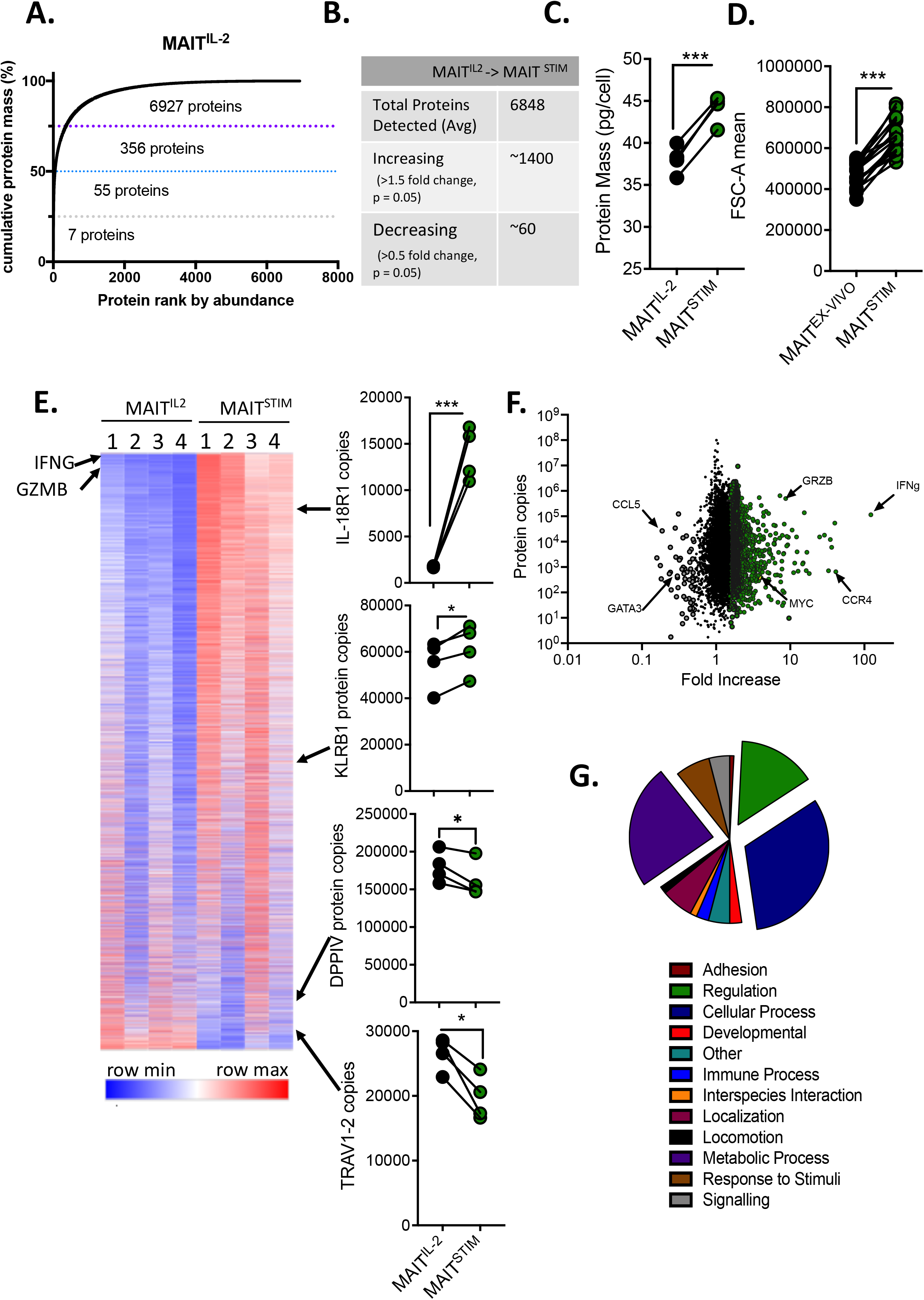
Remodelling of MAIT cell proteome after activation. (A) Proteins from IL-2 expanded MAIT (MAIT^IL-2^) cells were ranked by mass contribution and the mean cumulative protein mass was plotted against protein rank. (B) Table outlining the total number of proteins detected in MAIT^IL-2^ cells and their change upon stimulation (MAIT^STIM^; 18 hours with antiCD3/CD28 beads and 50ng/ml IL-12 & IL-18) using quantitative proteomics (n=4 per group). (C) Protein mass in MAIT^IL-2^ or MAIT^STIM^. (D) Scatter plot showing ex-vivo MAIT cell size (Forward scatter) basally or stimulated (as above). (E) Heatmap of the proteome of MAIT^IL-2^ and MAIT^STIM^ cells (as above). Relative protein abundance is graded from blue (low) to red (high) per row. Key MAIT cell proteins (IL-18R1, KLRB1, DPPIV & TRAV1-2) are represented in scatter graphs and their position in the heatmap is denoted by arrows). (F) Volcano plot displaying the MAIT^STIM^ cell proteome compared with MAIT^IL-2^ (Green circles represent proteins increased more than 1.5 fold over MAIT^IL-2^ cells, and grey dots represent proteins decreased more than 1.5 fold). (G) Pie chart proportional pathway analysis based on the MAIT^STIM^ cell proteome. * p<0.05, ** p<0.01 and *** p<0.001.

### MAIT cells upregulate expression of the transcription factor MYC and its downstream target pathways upon activation

*In silico* pathway analysis of a previously published RNA sequencing data-set by Lamichhane and colleagues^8^ revealed that *ex-vivo* activated MAIT cells (MAIT^EX-VIVO^ cells) upregulate a strong MYC signature (Figure 2A & Supplementary Figure 1A-B). In our proteomic dataset, we also found a robust increase in MYC and its target proteins in the activated human MAIT^IL-2^ cells (Figure 2B-C). Using western blotting and flow cytometry we investigated whether immune stimulation (TCR/CD28 and IL-12/IL-18) induced MYC expression in MAIT^EX-VIVO^ cells as well as in MAIT^IL-2^ cells (Figure 2D-G). To further verify the increased expression of MYC targets in activated MAIT cells we interrogated another published RNA sequencing data set by Hinks *et al*, which also demonstrated a robust increase in MYC target gene expression in activated MAIT cells (Figure 2H & Supplementary Figure 2).

**Figure 2.**
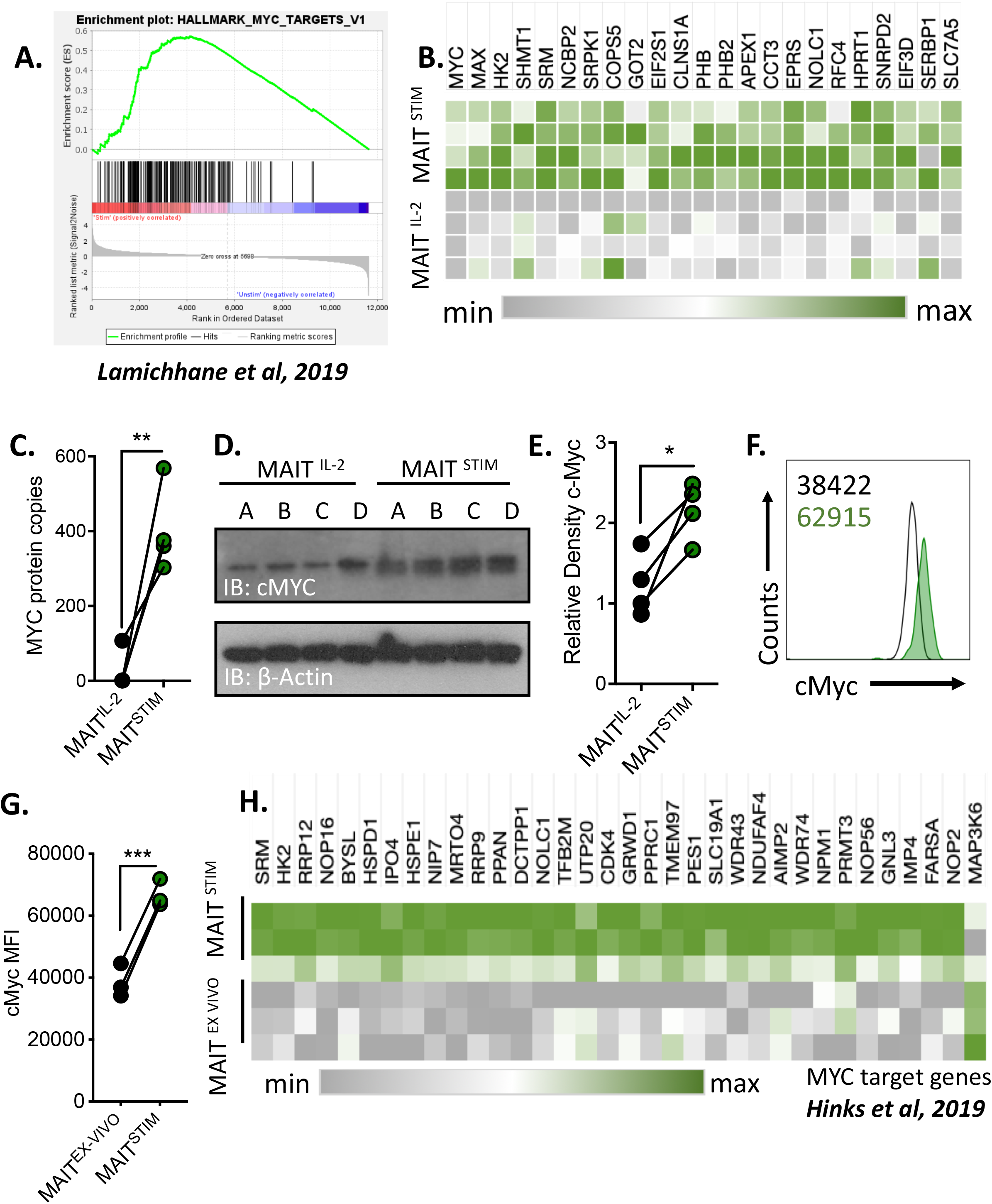
Activation of MYC and its target pathways in MAIT cells. (A) GSEA plot showing the increase in MYC target gene expression in activated MAIT cells. Data extrapolated from RNA sequence dataset published by Lamichhane *et al* on *ex vivo* MAIT cells stimulated via TCR. (B) Heatmap displaying expression levels of MYC and MYC-target in basal (MAIT^IL-2^) or stimulated (MAIT^STIM^; 18hr antiCD3/CD28 beads + 50ng/ml IL-12 & IL-18) cells as determined by quantitative proteomics (n=4 per group). Relative protein abundance is graded from grey (low) to green (high) per row. (C) Scatter plot showing MYC protein copies in basal (MAIT^IL-2^) or stimulated (MAIT^STIM^) cells. (D-E) Western blot and densitometry displaying MYC expression in MAIT^IL-2^ or stimulated (as above) MAIT^STIM^ cells. (F-G) Representative flow cytometry histogram (F) (Mean Fluorescence Intensity (MFI) values are shown within histogram) and MFI values (G) of MYC expression in *ex vivo* MAIT cells (black) or stimulated with 18hr antiCD3/CD28 beads + 50ng/ml IL-18 (green). (H) Heatmap displaying MYC target genes in *ex vivo* MAIT cells with or without TCR stimulation. Data extrapolated from RNA sequence dataset published by Hinks *et al*. * p<0.05, ** p<0.01 and *** p<0.001.

### MAIT cell proliferation is dependent on MYC

Previous studies have demonstrated that upon activation by TCR triggering with cognate antigen or bacterial infection MAIT cells can readily proliferate. To build on these observations we investigated the ability of MAIT^EX-VIVO^ cells to proliferate *in vitro* after stimulation with their cognate antigen 5-ARU-MG. Intriguingly, MAIT^EX-VIVO^ cells activated with their cognate antigen, 5-ARU-MG, fail to proliferate despite increased expression of activation markers such the Interleukin 2 (IL-2) receptor alpha chain CD25 (Figure 3A-C). This robust increase of CD25 expression was also noted on MAIT^IL-2^ cells after stimulation (Figure 3D). IL-2 is a well-established driver of T cell growth and proliferation^26^, so we investigated if the addition of IL-2 to 5-ARU-MG stimulation could drive MAIT^EX-VIVO^ cell proliferation. Addition of IL-2 resulted in significant MAIT cell growth and proliferation in response to 5-ARU-MG stimulation (Figure 3E-G). MYC expression is critical for conventional T cell blasting and proliferation in response to TCR signalling^18, 19^. We made use of the MYC inhibitor 10074-G5, which blocks binding of MAX to MYC and thereby inhibits transcriptional activity, and demonstrate that this increase in size is dependent upon MYC activity, and increase in CD25 expression also was dependent on MYC activity (Figure 3H-I). Furthermore, MYC activity is critical for MAIT cell proliferation in response to TCR and IL-2 (Figure 3J & Supplementary Figure 3).

**Figure 3.**
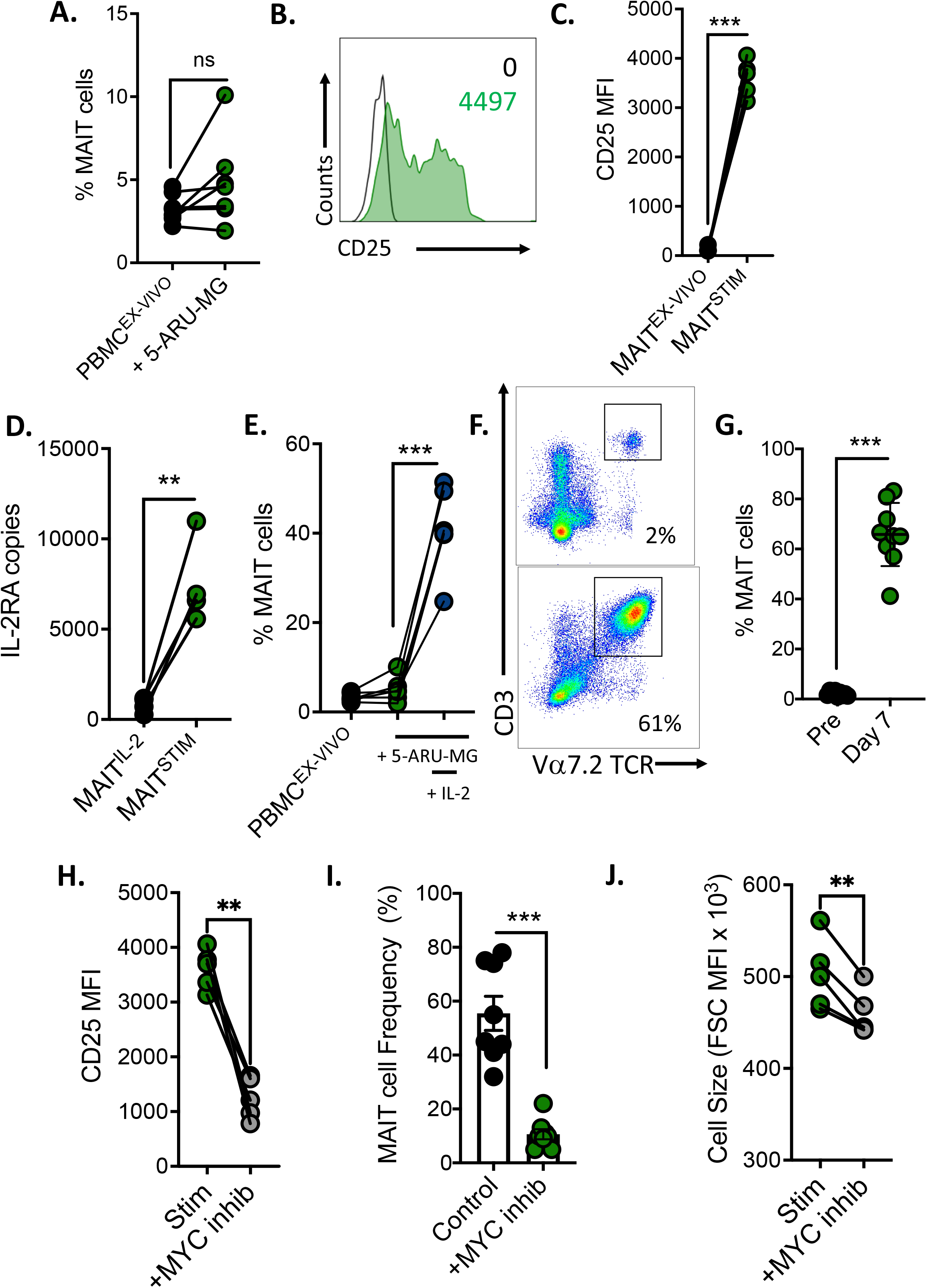
IL-2 driven proliferation of MAIT cells is dependent on MYC. (A) The frequency of MAIT cells (as a percentage of total T cells) after stimulation of PBMC (1×10^6^) with cognate antigen 5-ARU (1ug/ml) and Methylglyoxal (100μM) for 7 days (n=6). (B-C) Representative flow cytometry histogram and scatter plot showing CD25 (IL-2RA) expression on MAIT^ex vivo^ or stimulated through the TCR using antiCD3/CD28 beads/IL-12 & IL18 (MAIT^STIM^) for 18 hours (n=9). (D) Scatter plot showing CD25 (IL-2RA) protein copies in MAIT^IL-2^ cells basally or stimulated through the TCR using antiCD3/CD28 beads/IL-12 & IL18 (MAIT^STIM^) (n=4). (E) Line graph demonstrating frequency of MAIT cells (represented as percentage of T cells) after stimulation with 5-ARU (1ug/ml) and Methylglyoxal (100μM) in the absence or presence of IL-2 (35ng/ml) for 7 days (n=5). (F-G) Representative flow cytometry dot plot and scatter plot showing MAIT cell frequencies after 7 days stimulation with 5-ARU, Methylglyoxal and IL-2 (n=10). (H) MFI of CD25 expression on MAIT cells after 18 hours stimulation (as in C) in the absence or presence of the MYC specific inhibitor (10074-G5, 10uM) (n=5). (I) MAIT^IL-2^ cell frequency after 7 days expansion (as in E) in the absence of presence of the MYC specific inhibitor 10074-G5 (10uM). (J) Forward scatter MFI of MAIT cells after stimulation (as in C) in the absence of presence of the MYC inhibitor 10074-G5. * p<0.05, ** p<0.01 and *** p<0.001.

### The MYC-controlled amino acid transporter SLC7A5 is required for MAIT cell proliferation

Proliferation is a metabolically intense process, requiring significant energy and *de novo* generation of biosynthetic intermediates, so we next investigated metabolic processes under the control of MYC. A recent study by Marchingo and colleagues demonstrated that, in conventional T cells, TCR-regulated MYC expression induces the expression of critically important amino acid transporters, including SLC7A5^19^. SLC7A5 and its heavy chain chaperone, CD98 (aka SLC3A2) form the heterodimeric large neutral amino acid transporter (LAT-1). We made use of a flow cytometry-based assay, which measures uptake of the fluorescent SLC7A5 substrate kynurenine, to monitor uptake through SLC7A5 in MAIT cells. *Ex vivo* MAIT cells have low levels of kynurenine uptake, however upon TCR/cytokine stimulation, MAIT^EX-VIVO^ cells rapidly increase transport through SLC7A5 (Figure 4D). This correlates with increased expression of the heavy chain chaperone CD98 (Figure 4C). We next asked if stimulation of MAIT^IL-2^ cells also regulated SLC7A5 expression. Interrogation of the MAIT cell proteomes show that SLC7A5 expression increases from 20000 copies per cell in MAIT^IL-2^ cells to 40000 copies per cell in MAIT^IL-2+STIM^ cells (Figure 4A) which correlates with increased mRNA expression in MAIT^IL-2+STIM^ compared with MAIT^IL-2^ cells (Figure 4B). Together these data show that that upon activation both MAIT^EX-VIVO^ and MAIT^IL-2^ cells increase the expression of SLC7A5 protein, *SLC7A5* mRNA and CD98 surface expression (Figure 4A-C). We used MYC inhibitors to investigate whether the TCR/cytokine driven increase in LAT-1 expression was dependent upon MYC activity. Inhibition of MYC resulted in reduced SLC7A5 mRNA expression as well as reduced CD98 expression in activated MAIT cells (Figure 4E-F). Next, we investigated if loss of amino acid transport via LAT-1 impacted MAIT cell proliferation. Using the competitive substrate BCH to block LAT-1 activity, we show that, MAIT cell proliferation is limited by blocking uptake through LAT-1. Therefore, SLC7A5 activity during TCR/IL-2 activation is critical for MAIT^EX-VIVO^ cell proliferation (Figure 4G-I).

**Figure 4.**
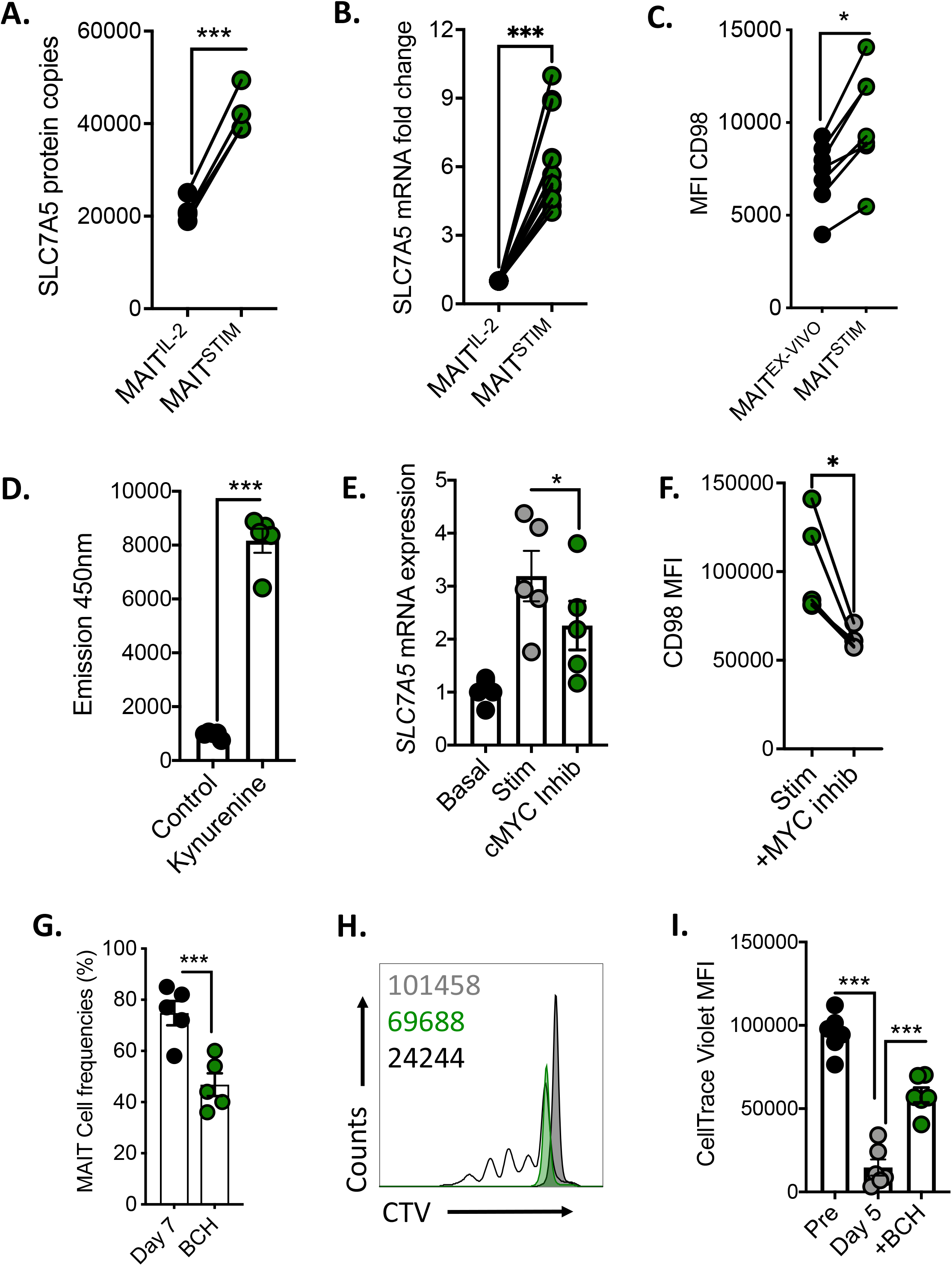
SLC7A5 facilitates MAIT cell proliferation and is dependent on MYC. (A) Mean protein copy number per cell of SLC7A5 in MAIT^IL-2^ cells and in cells stimulated for 18hrs (MAIT^STIM^; antiCD3/CD28 beads/IL-12 & IL18) (n=4). (B) Scatter plot showing SLC7A5 mRNA expression in MAIT^IL-2^ cells or MAIT^STIM^ (as in A) (n=12). (C) Flow cytometry MFI data of CD98 expression on ex-vivo MAIT cells or stimulated for 18 hrs (as in A)(n=7). (D) Flow cytometry data of uptake of kynurenine into ex vivo MAIT cells (n=5). (E) SLC7A5 mRNA expression in MAIT^IL-2^ cells and MAIT^STIM^ (as in A) in the absence or presence of the MYC specific inhibitor 10074-G5. (F) Flow cytometry MFI expression of CD98 on ex-vivo MAIT cells after stimulation (as in A) in the absence or presence of the MYC specific inhibitor 10074-G5 (10uM) (n=5). (G) The frequency of MAIT cells after 7 days expansion with 5-ARU (1ug/ml) and Methylglyoxal (100μM) in the absence or presence of IL-2 (35ng/ml)) in the absence of presence of the SLC7A5 inhibitor BCH (n=5). (H-I) Representative flow cytometry histogram and scatter plot showing cell trace violet (CTV) MFI in MAIT cells after 5 days stimulation (as in G) in the absence of presence of BCH (n=7). * p<0.05, ** p<0.01 and *** p<0.001.

### Glucose metabolism supports MAIT cell proliferation

MYC has been highlighted as a master regulator of glycolysis, therefore we investigated if glucose metabolism was required for MAIT cell proliferation^18, 19^. Previously we reported that MAIT cells increased glycolytic metabolism upon activation, and in agreement with our previous findings we noted increased expression of the glycolytic enzymes (hexokinase-II (HKII) and lactate dehydrogenase (LDH)) in both MAIT^EX-VIVO^ and MAIT^IL-2^ cells after 18 hours of stimulation (Figure 5A-D). Inhibition of MYC resulted in diminished HKII mRNA expression in activated MAIT^EX-VIVO^ cells (Figure 5E). To investigate if glucose metabolism is required for MAIT cell proliferation, we first utilized the glycolytic inhibitor 2-deoxy-D-glucose (2DG) and show inhibition of proliferation (Figure 5F-H). To support this finding, we limited the glucose availability in our culture media from 10mM to 1mM and show reduced MAIT cell proliferation (Figure 5I-J). Finally, we switched the carbon source in our culture media from glucose to galactose, which results in substantially lower rates of glycolysis and again show reduced MAIT cell proliferation (Figure 5K-L).

**Figure 5.**
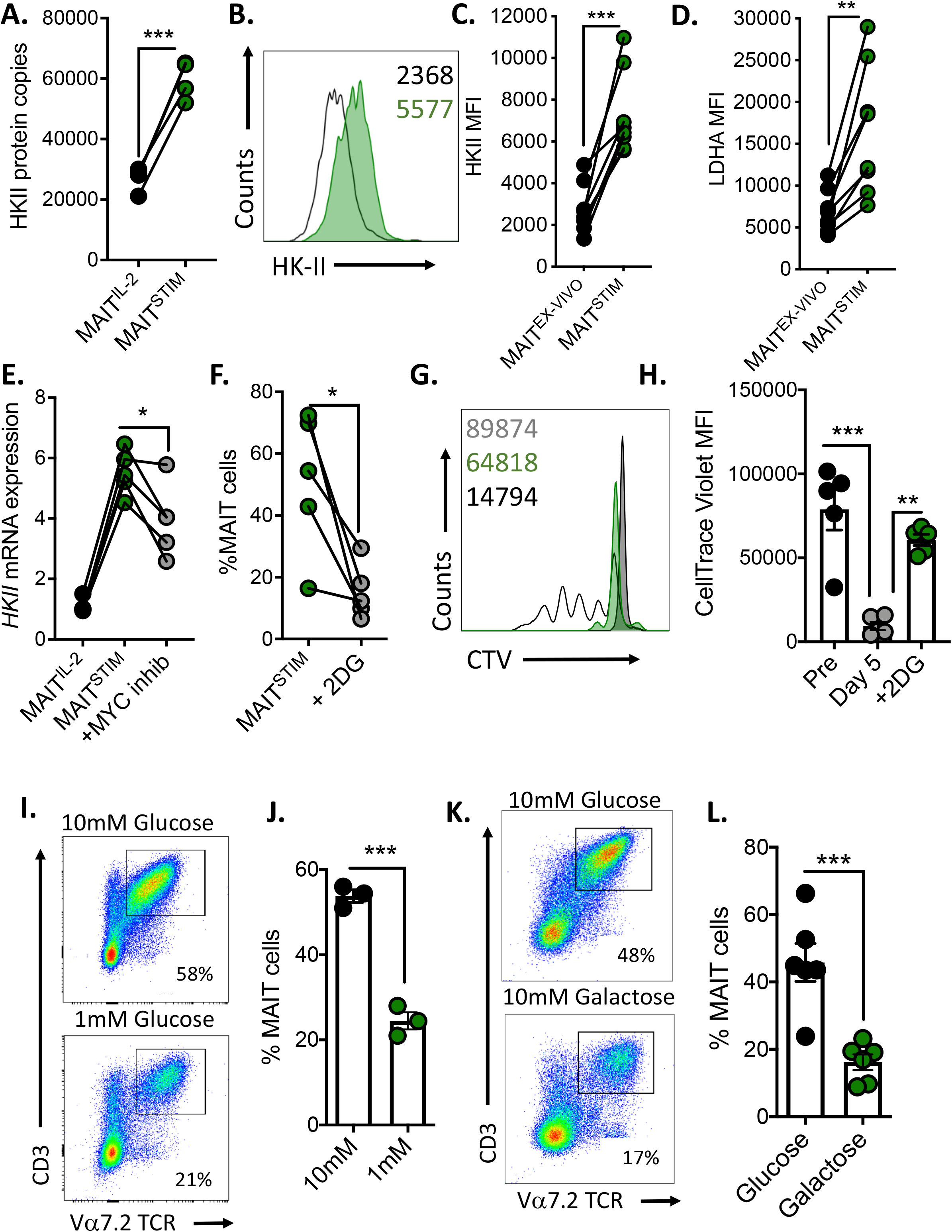
Glucose metabolism facilitates MAIT cell proliferation and is dependent on MYC. (A) Mean protein copy number per cell of Hexokinase-II (HKII) in MAIT^IL-2^ and in MAIT^IL2+STIM^ (stimulated for 18 hours with antiCD3/CD28 beads and IL-18 (50ng/ml) (n=4). (B-C) Representative flow cytometry histogram and MFI data of HKII in MAIT ^ex-vivo^ cells basally or stimulated (18 hours with antiCD3/CD28 beads and IL-18 (50ng/ml); MAIT^STIM^)(n=8). (D) Flow cytometry MFI data of LDHA expression in MAIT ^ex-vivo^ and MAIT^STIM^ cells (as in C) (n=8). (E) HKII mRNA expression in MAIT^IL-2^ and in and in MAIT^IL2+STIM^ cells (as in A) in the absence of presence of the MYC specific inhibitor 10074-G5 (n=5). (F) The frequency of MAIT cells after 7 days expansion from PBMC (1×10^6^) using cognate antigen 5-ARU (1ug/ml), Methylglyoxal (100μM) and IL-2 (35ng/ml) in the absence of presence of the glycolysis inhibitor 2DG (2mM) (n=5). (G-H) Representative flow cytometry histogram and scatter plot showing cell trace violet (CTV) MFI in MAIT cells after 5 days stimulation (as in F) in the absence of presence of 2DG (n=5). (I-J) Representative flow cytometry dot plot (I) and MAIT cell frequencies (J) after 7 days stimulation (as in F) in media containing either 10mM or 1mM glucose (n=3). (K-L) Representative flow cytometry dot plot (K) and MAIT cell frequencies (L) after 7 days stimulation (as in F) in media containing either 10mM glucose or 10mM galactose (n=6). * p<0.05, ** p<0.01 and *** p<0.001.

### MAIT cells from people with obesity display defective MYC and fail to proliferate

Previous work from our lab and others has demonstrated altered MAIT cell frequencies and cytokine production in people with obesity (PWO)^10, 27, 28^. We investigated the proliferative capacity of MAIT cells from PWO in response to antigenic (5-ARU-MG) and cytokine (IL-2) stimulation. We show that MAIT cells from PWO display defective proliferation in response to immune stimulation when compared to healthy controls (Figure 6A-B). These data provide further evidence that MAIT cells are functional impacted by obesity. Having highlighted the critical importance of MYC for MAIT cell proliferative responses, we assessed MYC expression in cohorts of PWO and healthy controls, and demonstrate diminished induction of MYC expression in MAIT^EX-VIVO^ cells in response to immune activation from PWO (Figure 6C-D). Targeted analysis of a previously published RNA sequencing data set from our group on MAIT cells isolated from PWO and healthy controls^22^, shows reduced expression of MYC target gene expression in MAIT cells from PWO (Figure 6E). Optimal CD25 surface protein expression was regulated by MYC (Figure 3H), so we investigated CD25 expression on MAIT cells from PWO, and see diminished CD25 levels in response to TCR and cytokine activation (Figure 6F). Having identified that SLC7A5, a direct transcriptional target of MYC, is critical for MAIT cell proliferation in response to immune stimulation, we investigated SLC7A5 expression on MAIT cells from PWO. We demonstrate that upon activation MAIT cells from PWO express significantly less SLC7A5 mRNA and CD98, resulting in reduced amino acid transport, a process critical for proliferation (Figure 6G-I).

**Figure 6.**
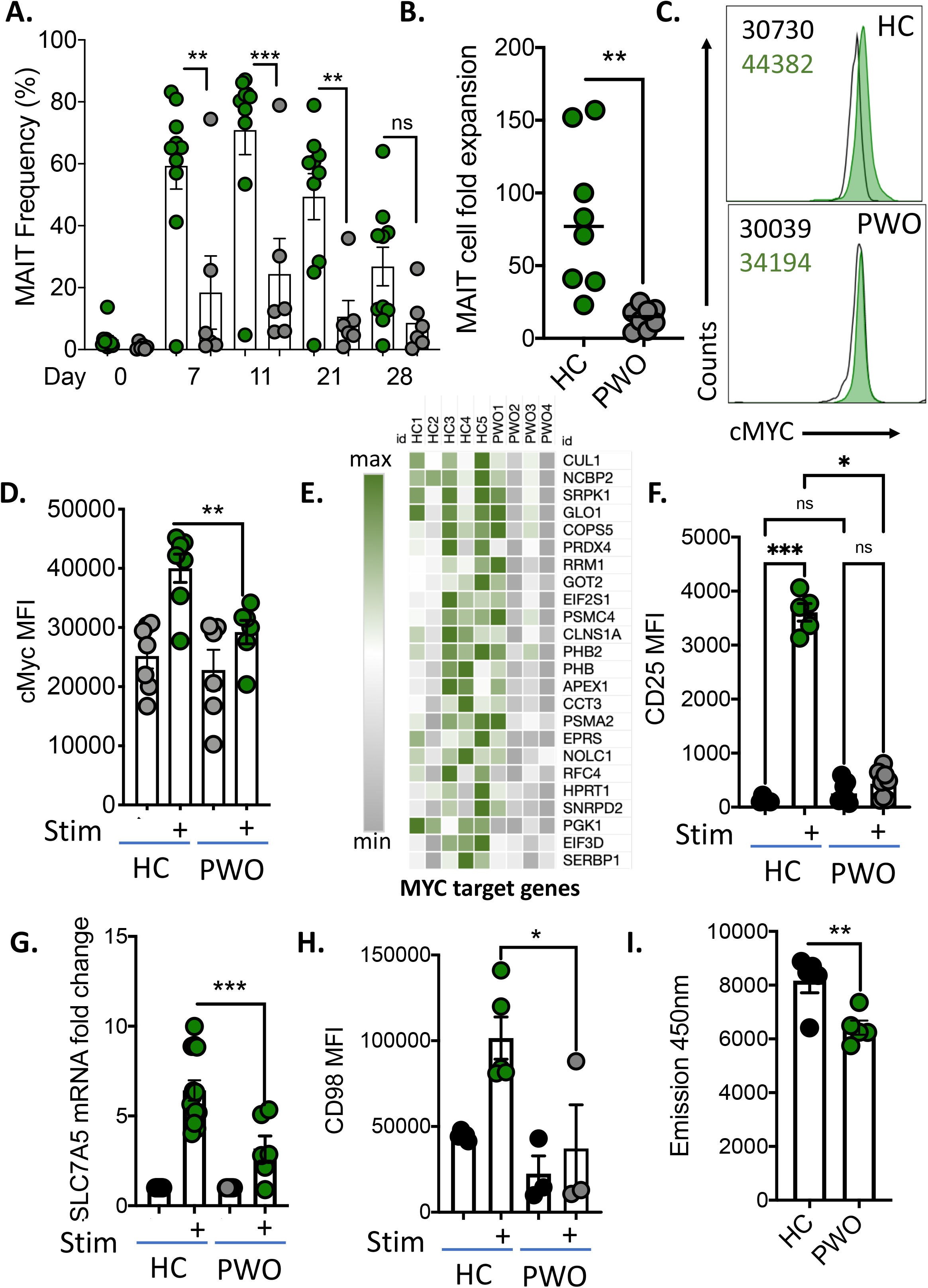
MYC is defective in obesity resulting in diminished MAIT cell proliferation. (A) The frequency of MAIT cells after 0-28 days expansion after 18 hour stimulation with 5-ARU (1ug/ml), Methylglyoxal (100μM) and maintenance in IL-2 (35ng/ml) in either healthy controls (green circles) or people with obesity (PWO) (grey circles) (n=10/group). (B) Fold expansion of MAIT cells from healthy controls or people with obesity after 7 days stimulation (with 5-ARU (1ug/ml), Methylglyoxal (100μM) and IL-2 (35ng/ml)) (n=8). (C-D) Representative flow cytometry histogram (C) and MFI data (D) of MYC expression in ex vivo MAIT cells from healthy controls (top panel,C) or PWO (bottom panel,C) stimulated with antiCD3/CD28 beads and IL-18 (50ng/ml) for 18 hours (n=6-7/group). (E) Heatmap displaying MYC target genes in ex-vivo MAIT cells from health controls or PWO (n=4-5/group). (F) CD25 expression (MFI) on ex-vivo MAIT cells or stimulated MAIT cells (as in C) from either healthy controls or PWO (n=5-8/group). (G) mRNA expression of *Slc7a5* in MAIT^IL-2^ cells or MAIT^IL-2^ stimulated (as in C) from either healthy controls or PWO (n=6-10/group). (H) Scatter plot showing CD98 expression (MFI) on ex-vivo MAIT cells basally or stimulated (as in C) from either healthy controls or PWO (n=3-4/group). (I) Scatter plot showing Kynurenine uptake into MAIT cells from either healthy controls or PWO (n=5/group). * p<0.05, ** p<0.01 and *** p<0.001.

## Discussion

MAIT cells are a subset of unconventional T cells which are abundant in human blood and tissues, including liver and adipose tissue^29, 30^. MAIT cells are capable of rapidly responding to stimulation, producing cytokines, lytic molecules and proliferating^29^. Due to their potent effector functions and abundance, MAIT cells have been shown to play an important role in the host defence against pathogens and malignancies^1, 31^, and are now under investigation as a potential immunotherapeutic agent^16, 32^. However, the molecular regulation of MAIT cell effector functions are still emerging. In the current study we demonstrate for the first time that MAIT cell proliferation is dependent on the transcription factor MYC. We show that upon activation MAIT cells robustly increase MYC target proteins including, SLC7A5 and HKII, which are required for MAIT cell proliferation. Finally, we show for the first time that MYC expression and activity is defective in people with obesity, underpinning diminished MAIT cell proliferation, which may increase host susceptibility to infection and malignancies.

Quantitative mass spectrometry allows a high dimensional analysis of the proteome and activation induced remodelling^19, 20, 21^. Using this approach, we were able to identify >6800 proteins in both resting IL-2 expanded MAIT cells and stimulated IL-2 expanded MAIT cells, significantly more than the identification of 3000-5500 protein previously published in MAIT cells^33, 34^. TCR and cytokine stimulation was associated with significant remodelling of the MAIT cell proteome with significant increases in protein content and cell size. Pathway analysis of both our proteomic dataset and previously published RNA sequencing datasets revealed MYC as one of the most upregulated pathways in MAIT cells, however its role remains unclear. MYC is a critically important transcription factor in conventional T cells, and acts as a metabolic master regulator^18, 19, 21^. Experiments using high-dimensional quantitative mass spectrometry on CD4 and CD8 T cells from wild type and MYC deficient mice revealed how MYC controls TCR driven cell growth and metabolism^18, 19^. We show for the first time that MYC is a critical regulator of MAIT cell proliferation. MAIT cell proliferation was triggered in a two-step process, activation with the cognate MAIT cell antigen 5-ARU did not drive MAIT cell proliferation, but did trigger the expression of the high affinity IL-2 receptor CD25. IL-2 is well established as a potent driver of T cell proliferation, including MAIT cells^16^, and here we confirm these observations, with the addition of IL-2 resulting in robust proliferation of MAIT cells. A similar two-step process has previously been reported in conventional CD8 T cells, with IL-2 not required for the initiation of proliferation but critical for sustaining proliferation^35^. We also demonstrate that increased expression of CD25 by activated MAIT cells was dependent on MYC, which aligns with previously published data which showed that in MYC deficient conventional T cells, the expression of CD25 was not increased upon stimulation^19^.

MYC signalling is also essential for the upregulation of amino acid transporters on T cells including the most abundantly expressed SLC7A5^19^. We have previously reported that MAIT cells express SLC7A5^22^, and demonstrate here that inhibition of MYC results in diminished SLC7A5 and CD98 expression. Amino acid transport via SLC7A5/CD98 is critical for sustained expression of MYC, suggesting a positive feed-forward loop^19^. Using the SLC7A5 inhibitor BCH we show that amino acid transport through SLC7A5 is critical for MAIT cell proliferation, highlighting for the first time a MYC-SLC7A5 axis in activated MAIT cells.

MYC also directs a glycolytic metabolism programme in conventional T cells^18, 19, 21^, which is well established as a critical process for immune growth, activation and effector functions^36^. We and others have previously reported that glycolytic metabolism is critical for MAIT cell cytokine and lytic molecule production ^22, 27, 37^ but the factors regulating MAIT cell metabolism and the requirements for MAIT cell proliferation are unknown. Herein we show that MAIT cells increase the expression of key glycolytic enzymes (e.g. HKII and LDH), supporting MAIT cell engagement of glycolytic metabolism upon activation. Furthermore, inhibition of MYC resulted in diminished HKII expression, confirming that as in conventional T cells MYC is a key metabolic regulator. In conventional CD8 T cells, glucose metabolism is also critical for cell growth and proliferation^38^. We show using a series of approaches including inhibition of glycolysis with low dose 2-DG that glucose metabolism is critical for MAIT cell proliferation. Limiting glycolysis by acutely reducing glucose availability or substituting glucose with galactose also significantly impeded MAIT cell proliferation, providing evidence for glucose availability being a rate limiting step.

MAIT cell cytokine production has been reported as defective in numerous human diseases, including cancer, obesity and more recently COVID-19^31, 39, 40^. Here we show for the first time that MAIT cell proliferation is defect in people with obesity, which may link with the reports of defective host protection in obesity. Having demonstrated the importance of MYC and its target pathways for MAIT cell proliferation, we investigated the impact of obesity on MYC for the first time, and show reduced MYC activity in MAIT cells from people with obesity. Furthermore, expression of key MYC targets like SLC7A5 were also defective in MAIT cells isolated from people with obesity. We have previously demonstrated that obesity associated defects in glucose metabolism in both MAIT cells and NK cells underpins defective cytokine production^22, 41, 42^. The identification of defective MYC in obesity helps to further understand these observations.

In conclusion, we identify MYC as a critical regulator of MAIT cell metabolism. We demonstrate that a MYC-SLC7A5-glycolysis axis is critical for MAIT cell proliferation and is defective in obesity. However, this data extends beyond obesity and provides important insight into the molecular and metabolic regulation of MAIT cell proliferation which will have particular relevance for the potential use of MAIT cells for immunotherapy^16^.

## Materials & methods

### Study cohorts & ethical approval

A total cohort of 50 adults (25obese/25 non-obese) were recruited. Inclusion criteria included ability to give informed consent, 18-65 years of age and a BMI<28 for the non-obese control cohort and BMI >30 for obese cohort. Exclusion criteria for both cohorts included current or recent (<2 weeks) infection, current smoker, use of anti-inflammatory medications including GLP-1 analogue therapies. Ethical approval was obtained from both St Vincent’s University Medical Ethics Committee and Maynooth University Ethics Committee

### Preparation of peripheral blood mononuclear cells (PBMC) and flow cytometric analysis

PBMC samples were isolated by density centrifugation over ficoll from fresh peripheral blood samples. MAIT cell staining was performed using specific surface monoclonal antibodies (All Miltenyi Biotec) namely; CD3, CD161, CD8 and TCRVα7.2. Cell populations were acquired using a Attune NXT flow cytometer and analysed using FlowJo software (Treestar). Results are expressed as a percentage of the parent population as indicated and determined using flow minus-1 (FMO) and unstained controls.

### MAIT cell proliferation analysis

Fresh PBMC (1 × 10^6^ /ml) were stimulated for 18 hours with 1μg/mL of 5-ARU and 100μM of Methylglyoxal, in the absence or presence of specific metabolic inhibitors (2DG (2mM), 10074-G5(10μM), iBET762(10μM) or BCH(50mM)). After 18 hours, media was replaced with fresh culture media containing IL-2 (35ng/ml). Cultures were maintained for up to 28 days, replacing media with fresh culture media containing IL-2 every 3 days. MAIT cell proliferation was determined by either flow cytometric analysis of MAIT cell frequencies or cell trace violet proliferation assays.

### MAIT cell proteomic sample preparation

Purified IL-2 expanded MAIT cells (MAIT^IL-2^) were stimulated for 18 hours with anti-CD3/CD28 Dynabeads (Thermofisher), IL-12 (50ng/ml) and IL-18 (50ng/ml) for 18 hours. Cell pellets were lysed at room temperature in 4% SDS, 50mM TEAB pH8.5, 10mM TCEP under agitation for 5 mins, the boiled for 5 mins and sonicated with a BioRuptor (30 seconds on, 30 seconds off x 30 cycles). Protein concentration was determined using EZQ protein quantification kit (Invitrogen) according to the manufacturer’s protocol. Lysates were alkylated with 20mM iodoacetamide for 1 hours at room temperature in the dark. The samples were then processed using S-Trap mini columns (Protifi). 12% aqueous phosphoric acid was added at 1:10 to each sample for a final concentration of ~1.2% phosphoric acid. Samples were transferred to 5ml lo bind Eppendorf tubes (Eppendorf). 3200μl of S-Trap binding buffer (100mM TEAB - pH 7.1 adjusted using phosphoric acid, 90% MeOH) was added to each sample. 650μl of each sample was loaded onto an S-Trap column and spun at 4,000 g for 30 s or until all SDS lysate/S-Trap buffer had passed through the S-Trap column. Loading and spinning of columns was repeated until all the lysate was run through the column. Columns were transferred to fresh 2ml collection tubes. 125μl of digestion buffer (50mM ammonium bicarbonate in HPLC water) containing 1:20 trypsin was added onto each column. Samples were spun at 4000g for 30 seconds and any solution that passed through was returned to the top of the column. Tubes were incubated for 2 hrs at 47°C. 80μL of digestion buffer containing trypsin was added to each column. Samples were centrifuged at 1,000 g for 60 sec and the peptide elution kept. 80μL of 0.2% aqueous formic acid was added to the S-Trap protein-trapping matrix and spin through at 1,000 g for 60 sec into the same collection tube. 80μL of 50% aqueous acetonitrile containing 0.2% formic acid was added and samples spun at 4000g for 60 seconds for a final elution.

### Data independent acquisition (DIA) mass spectrometry acquisition

An equivalent of 1.5μg peptides were injected onto a nanoscale C18 reverse-phase chromatography column coupled to an UltiMate 3000 RSLC nano, HPLC system (Thermo Fisher) and an Orbitrap Exploris 480 Mass Spectrometer (Thermo Fisher). For liquid chromatography the following buffers were used: buffer A (0.1% formic acid in Milli-Q water (v/v)) and buffer B (80% acetonitrile and 0.1% formic acid in Milli-Q water (v/v). Samples were loaded at 10μL/min onto a trap column (100 μm × 2 cm, PepMap nanoViper C18 column, 5μm, 100 Å, Thermo Scientific) equilibrated in 0.1% trifluoroacetic acid (TFA).

The trap column was washed for 3 min at the same flow rate with 0.1% TFA then switched in-line with a Thermo Scientific, resolving C18 column (75 μm × 50 cm, PepMap RSLC C18 column, 2 μm, 100 Å). Peptides were eluted from the column at a constant flow rate of 300nl/min with a linear gradient from 3% buffer B to 6% buffer B in 5 min, then from 6% buffer B to 35% buffer B in 115 min, and finally to 80% buffer B within 7 min. The column was then washed with 80% buffer B for 4 min and re-equilibrated in 3% buffer B for 15 min. Two blanks were run between each sample to reduce carry-over. The column was kept at a constant temperature of 50°C.

The data was acquired using an easy spray source operated in positive mode with spray voltage at 2.6 kV, and the ion transfer tube temperature at 250°C. The MS was operated in DIA mode. A scan cycle comprised a full MS scan (m/z range from 350-1650), with RF lens at 40%, AGC target set to custom, normalised AGC target at 300%, maximum injection time mode set to custom, maximum injection time at 20 ms, microscan set to 1 and source fragmentation disabled. MS survey scan was followed by MS/MS DIA scan events using the following parameters: multiplex ions set to false, collision energy mode set to stepped, collision energy type set to normalized, HCD collision energies set to 25.5, 27 and 30%, orbitrap resolution 30000, first mass 200, RF lens 40%, AGC target set to custom, normalized AGC target 3000%, microscan set to 1 and maximum injection time 55 ms. Data for both MS scan and MS/MS DIA scan events were acquired in profile mode. The method used for the DIA mass spectrometry was based on the paper by Muntel *et al*. (2019).

### DIA data quantification and analysis

Quantification of reporter ions was completed using Spectronaut (VX, Biognosys; Spectronaut 14.10.201222.47784) in library-free (directDIA) mode. Minimum peptide length was set to 7 and maximum passport length was set to 52, with a maximum of 2 missed cleavages. MS1 and MS2 mass tolerance strategy and XIC IM and RT extraction windows were set to dynamic, all with a correction factor of 1. Trypsin was specified as the digestive enzyme used. The false discovery rate at the precursor ion level and protein level was set at 1% (protein and precursor Q-value cut-off). The max number of variable modifications was set to 5, with protein N-terminal acetylation, methionine oxidation and glutamine and asparagine deamidation set as variable modifications. Carbamidomethylation of cysteine residues was selected as a fixed modification. For calibration, the MS1 and MS2 mass tolerance strategy was set to the system default. Machine learning was set to across experiment, with a precursor PEP cut-off of 1, a protein Q-value cut-off of 0.01. Single-hit proteins were not excluded, with single-hits defined by stripped sequence. For quantification, amino acids were set to false and best N fragments per peptide was set to true, with a minimum of 3 and maximum of 300, and ion charge and type set to false. The major protein grouping was by protein group ID and the minor peptide grouping was set to stripped sequence. Modifications were set to none Major and minor group top N was set to false, with minor and major group quantities set to sum precursor quantity and sum peptide quantity respectively. Quantity at the MS-level was set to MS2 and quantity type to area. Proteotypicity filter was set to none, data filtering to Q-value and cross run normalisation was switched off. MS2 demultiplexing was disabled, the run limit for the directDIA library set to −1, with no profiling strategy or unify peptide peaks strategy. Data filtering and protein copy number quantification was performed in the Perseus software package, version 1.6.6.0. Proteins were quantified from unique peptides and razor peptides. Copy numbers were calculated using the proteomic ruler as described (Wisniewski *et al*., 2014). This method sets the summed peptide intensities of the histones to the number of histones in a typical diploid mouse cell. The ratio between the histone peptide intensity and summed peptide intensities of all other identified proteins is then used to estimate the protein copy number per cell for all the identified proteins. Further filtration of the data was completed to include proteins detected in at least ≥ 2 biological replicates.

### Cell trace violet proliferation assay

MAIT cell proliferation was also measured using a CellTrace™ Violet (CTV) proliferation kit (ThermoFisher) according to the manufacturer’s instructions. Briefly, CellTrace solution was prepared immediately prior to use to 5mM stock using DMSO. Next the dye for diluted to 5μM working concentration by adding appropriate amount of the stock solution into pre-warmed PBS. Isolated PBMC were stained at 10^6^ cells per mL of the PBS-dye solution. Cells were incubated for 20 min at room temperature, protected from light with circular agitation. Unbound dye was washed away with RPMI, and cell were incubated for at least 10 minutes to allow acetate hydrolysis of the dye. CTV stained cells were stimulated for 18 hours with 1μg/mL of 5-ARU and 100μM of methylglyoxal, in the absence or presence of specific metabolic inhibitors (2DG (2mM), 10074-G5(10uM), iBET762(10μM) or BCH (50mM)). After 18 hours, media was replaced with fresh culture media containing IL-2 (35ng/ml). Cultures were maintained for 5 days before analysis.

### MAIT cell metabolic analysis

Fresh PBMC (1 × 10^6^ /ml) or MAIT^IL-2^ cells were activated using Dynabeads, IL-12 (50ng/ml) and IL-18 (50ng/ml) for 18 hours. Cells were then labelled for extracellular markers, then fixed and permeabilized using the True-Nuclear Transcription Factor Buffer set (BioLegend) according to the manufactures instructions before intracellular staining with monoclonal antibodies specific for hexokinase II or lactate dehydrogenase (Abcam).

### MAIT cell BCH experiments

For SLC7A5 inhibition experiments, the concentration of amino acids in RPMI was diluted twofold using Hank’s balanced salt solution (HBSS; Invitrogen) in the presence or absence of BCH (50mM Sigma).

### PCR gene expression

mRNA was extracted from MAIT cells from healthy controls and PWO using EZNA Total RNA kit I (Omegabio-tek) according to the manufacturer’s protocol. Synthesis of cDNA was performed using qScript cDNA Synthesis kit (QuantaBio). Real time RT-qPCR was performed using PerfeCTa SYBR Green FastMix Reaction Mix (Green Fastmix, ROX™) (QuantaBio) and KiCqStart primer sets (Sigma).

### Western blotting analysis

Human MAIT^IL-2^ cells (2.5×10^6^) were cultured in 24-well plates stimulated with Dynabeads, IL-12 (50ng/ml) and IL-18 (50ng/ml) for 18 hours before harvesting for western blotting. Cells were lysed in NP-40 lysis buffer (50mM Tris-HCI, pH 7.4, containing 150 mM NaCl, 1% (w/v) IgePal, and complete protease inhibitor mixture (Roche)). Samples were resolved using SDS-PAGE and transferred to nitrocellulose membranes before analysis with anti-MYC (Cell Signalling) anti-β-Actin (Sigma) antibodies. Protein bands were visualised using enhanced chemiluminescence.

### In silico RNA sequencing analysis

Publicly available RNA sequencing data sets of MAIT cells were downloaded from Gene Expression Omnibus (GEO) accession number GSE123805 (Hinks *et al*., 2019) and National Center for Biotechnology Information (NCBI) Sequence Read Archive (SRA) accession number PRJNA559574 (Lamichhane *et al*., 2019). Raw read counts were downloaded from GEO for the Hinks *et al*., (2019) study and the data analysis pipeline to this point is detailed in the associated paper. Whereas raw sequencing data in format of FASTQ files were downloaded from the NCBI SRA for the Lamichhane *et al*., (2019) study and the following data analysis pipeline was implemented. The TrimGalore (v0.6.6) tool was used with Cutadapt (v1.15) and FastQC to apply quality and adapter trimming to FASTQ files. STAR (v2.7.9a) was used to align trimmed reads to the human genome (*Homo sapiens* high coverage assembly GRCh38 from the Genome Reference Consortium – GRCh38.p13) with the quantMode GeneCounts option to output read counts per gene. The Bioconductor package EdgeR (v3.28.1) was applied in R (v3.6.3) to identify statistically significant differentially expressed genes between patient groups. Biological and technical variation was accounted for by the negative binomial distribution of RNAseq count data using a generalization of the Poisson distribution model. The filterByExpr function was applied to remove lowly expressed genes. The data was normalized across library sizes, between samples using the trimmed mean of M-values (TMM) normalization method. Tagwise dispersions were estimated for the normalized dataset. *P-*values from multiple comparisons were corrected with the Benjamini-Hochberg method in EdgeR. For the comparisons between stimulations and controls, genes were considered significantly differentially expressed with an FDR adjusted *p*-value < 0.1. Variance Modeling at the Observational Level (VOOM) method within edgeR was used to output normalized read counts as LogCPM values. These were used to perform hierarchical clustering and to construct heatmaps in Gene Pattern’s online server (v3.9.11) and to perform Gene Set Enrichment Analysis (GSEA) (v4.1.0) with annotated HALLMARK genesets from the MSigDB (Molecular Signatures Database) collections (v6.2). Venn diagrams were constructed using InteractiVenn^43^.

### Statistics

Statistical analysis was completed using Graph Pad Prism 6 Software (USA). Data is expressed as SEM. We determined differences between two groups using student t-test and Mann Whitney U test where appropriate. Analysis across 3 or more groups was performed using ANOVA. Correlations were determined using linear regression models and expressed using Pearson or Spearman’s rank correlation coefficient, as appropriate. P values were expressed with significance set at <0.05.

### Contributors Statement

NKM, MMP, CR, RB, NAW, CDB and CON performed the experiments and carried out analysis and approved the final manuscript as submitted. NV and GSB provided MAIT cell reagents and aided in the design of MAIT cell activation experiments. CF performed the *in silico* RNA sequencing analysis. NWK and DOS recruited patient cohorts and provided relevant clinical data. AEH, NJ, DOS & LVS conceptualized and designed the study, analyzed the data, drafted the manuscript, and approved the final manuscript as submitted.

---

**Supplementary Figure 1.**
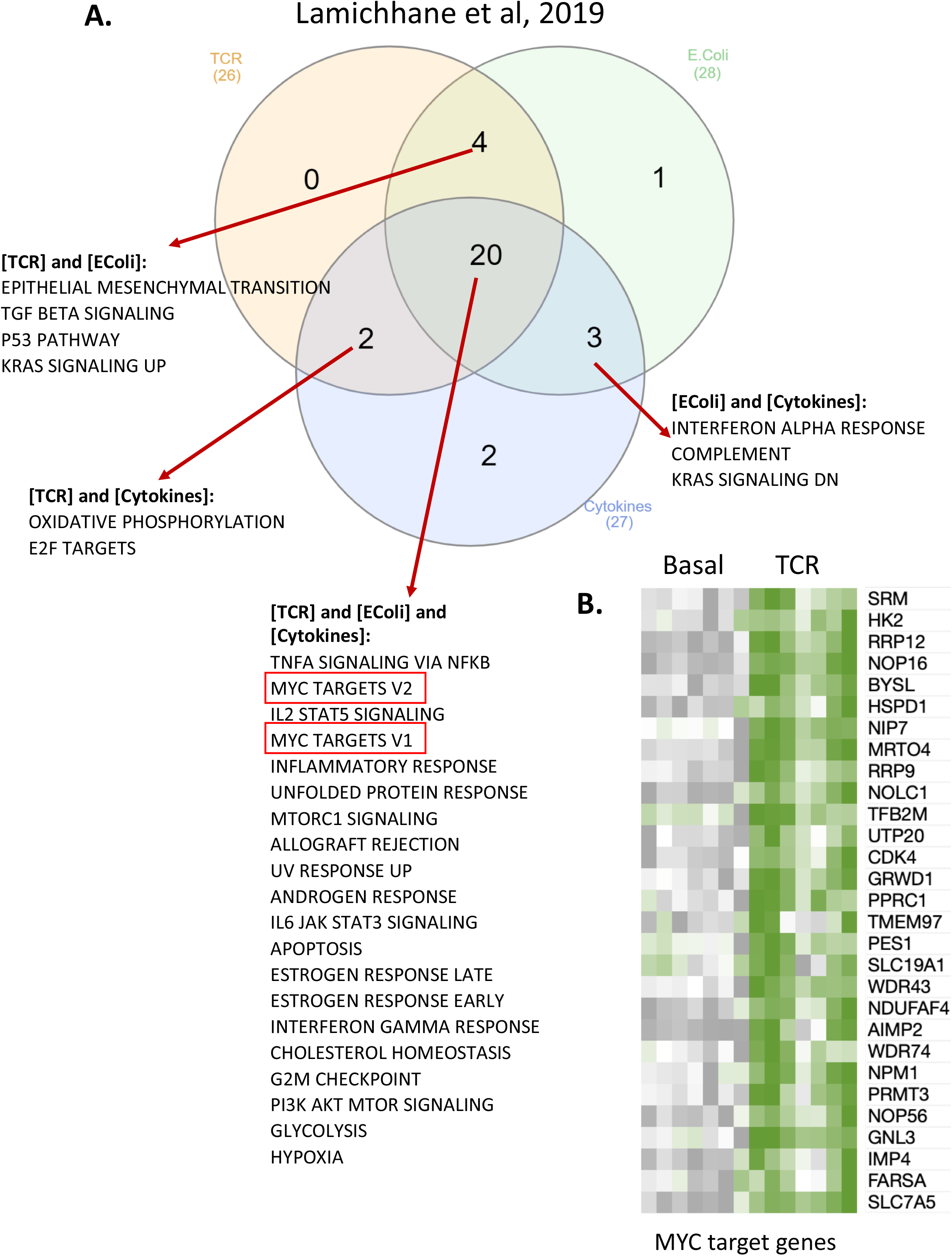
*In silico* transcriptomic analysis highlights upregulation of MYC target genes. (A) In silico transcriptomic analysis highlights the enrichment of common Gene Sets within differentially stimulated MAIT cells. (A) In silico analysis of data extrapolated from an RNA sequencing dataset published by Lamichhane et al (2019) on ex vivo MAIT cells stimulated via TCR, E.coli, cytokine or combinations. The Venn Diagram displays the number of significantly upregulated Hallmark Gene Sets following Gene Set Enrichment Analysis for each stimulation. (B) Heatmap of data from (A) displaying expression levels of significantly (FDR <0.1) differentially expressed MYC-target genes in unstimulated or TCR stimulated MAIT cells

**Supplementary Figure 2.**
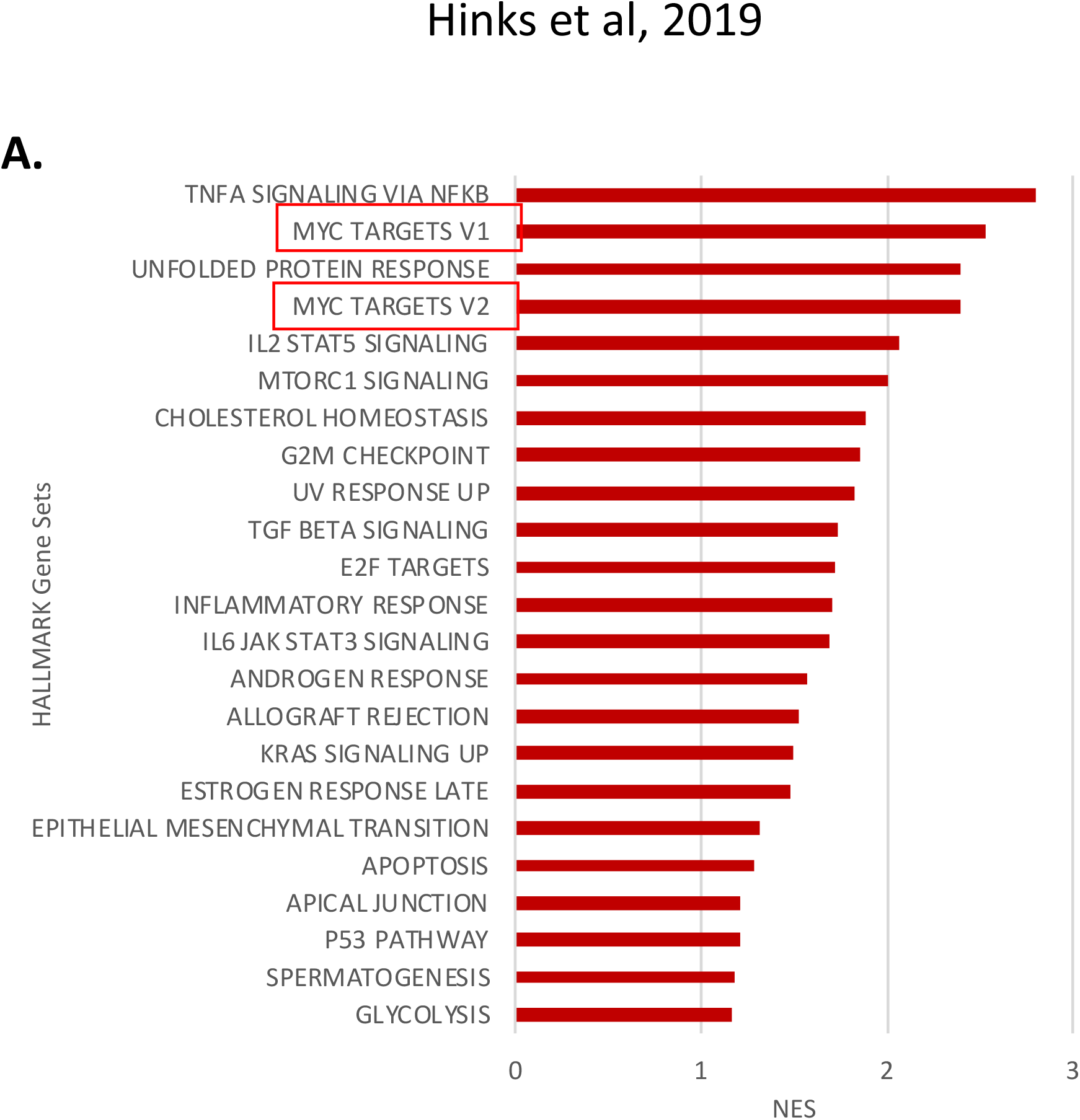
*In silico* transcriptomic analysis highlights upregulation of MYC target genes. (A) In silico transcriptomic analysis of MAIT cells stimulated through the TCR upregulates MYC target genes. (A) In silico analysis of data extrapolated from an RNA sequencing dataset published by Hinks et al (2019) showing significantly upregulated Hallmark Gene Sets following Gene Set Enrichment Analysis within TCR activated MAIT cells. Enriched Hallmark Gene Sets are ordered based on the Normalised Enrichment Score (NES).

**Supplementary Figure 3.**
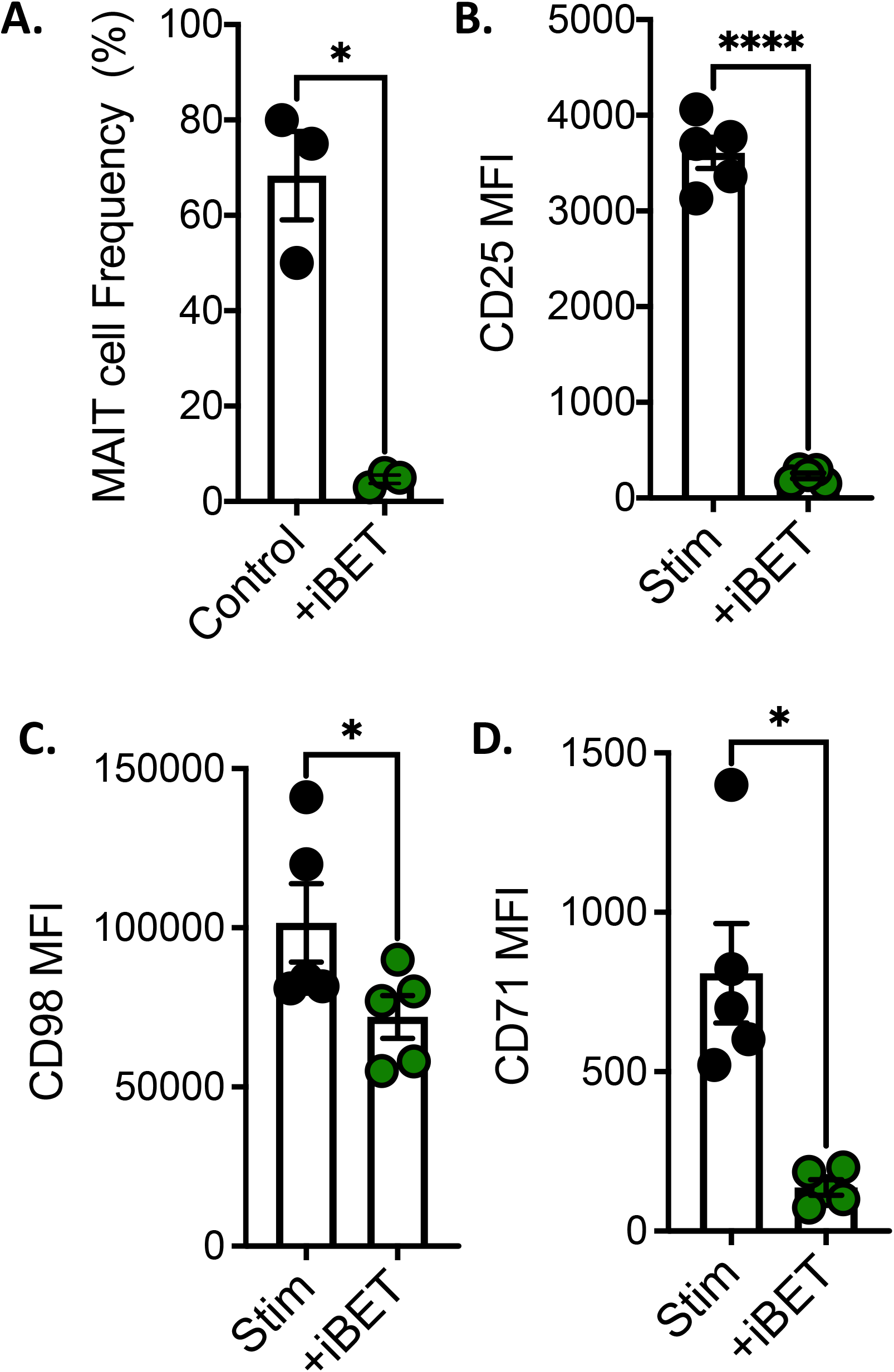
Inhibition of MYC with iBET762 limits MAIT cell proliferation and surface protein expression. (A) The frequency of MAIT cells (as a percentage of total T cells) after stimulation of PBMC (1×10^6^) with cognate antigen 5-ARU (1ug/ml) and Methylglyoxal (100μM) for 7 days in the absence or presence of iBET 762 for the first 18 hours (1⍰M) (n=3). (B-D) Scatter plots demonstrating the surface expression of CD25 (IL-2RA), CD98 or CD71 on MAIT^ex vivo^ stimulated through the TCR using antiCD3/CD28 beads/IL-12 & IL18 (MAIT^STIM^) for 18 hours in the absence or presence of iBET 762 (1⍰M) (n=5). * p<0.05 and **** p<0.001.

**Supplementary Figure 4.**
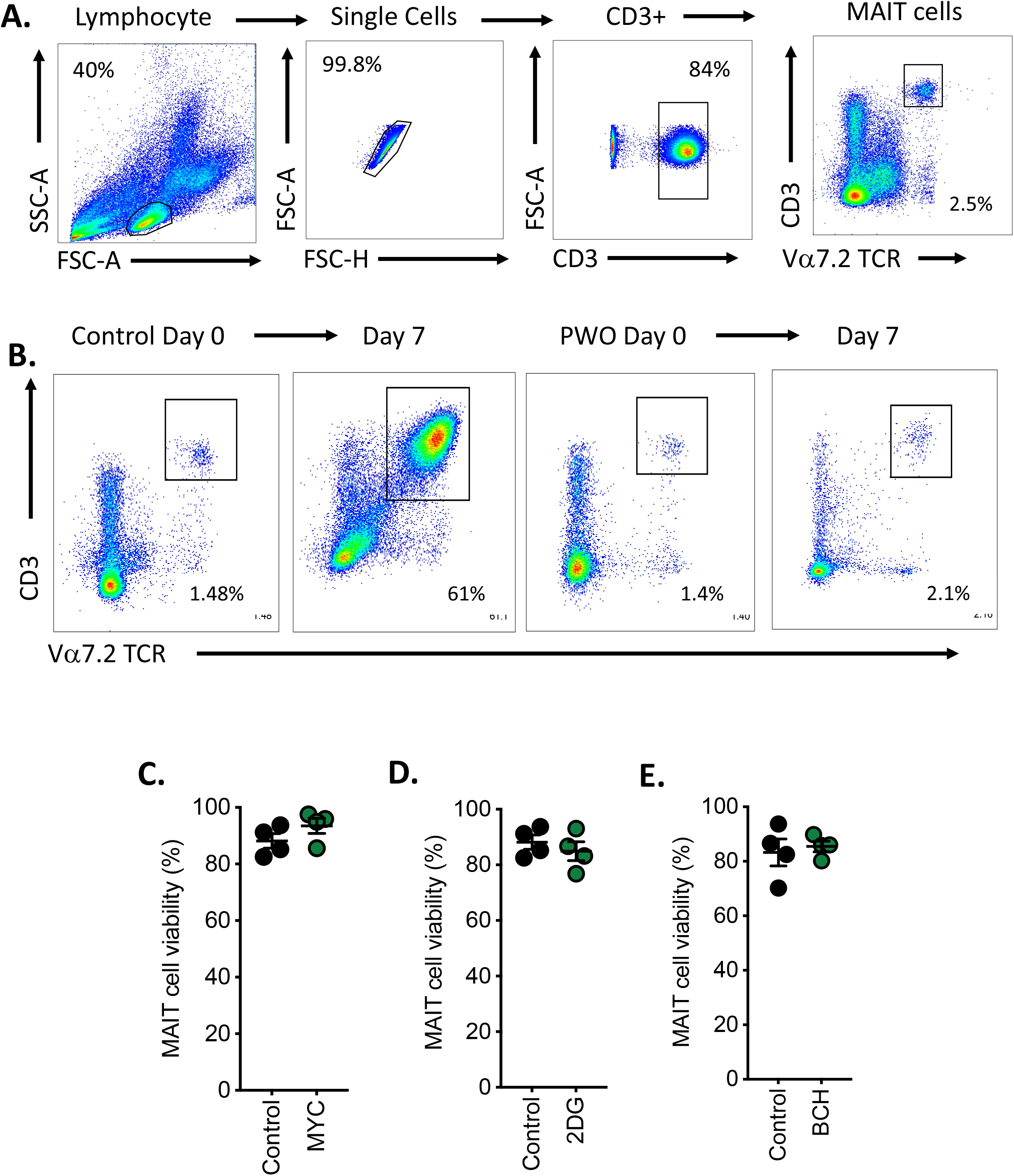
Gating strategy and cell viability. (A) Representative dot plots demonstrating the gating strategy for the identification of MAIT cells. (B) Representative dot plots from a healthy controls and a person with obesity (PWO) demonstrating the expansion of MAIT cells after stimulation of PBMC (1×10^6^) with cognate antigen 5-ARU (1ug/ml) and Methylglyoxal (100μM) for 7 days. (C-D) Scatter plots demonstrating the viability of MAIT cells (measured using zombie yellow viability dye) after 18 hours treatment with MYC specific inhibitor (10074-G5, 10uM, 2DG (2mM) or BCH (50mM) (n=3).

## Notes

**Funding Source**: This study is supported by the National Children’s Research Centre. NKM is supported by Health Research Board (ILP-POR-2019-110). CB is supported by a fellowship from Irish Research Council. Financial support for the Attune NxT was provided to Maynooth University Department of biology by Science Foundation Ireland (16/RI/3399).

**Conflict of Interest**: The authors declare no conflict of interest.

### Competing Interest Statement

The authors have declared no competing interest.

